# Unusual metabolism and hypervariation in the genome of a Gracilibacteria (BD1-5) from an oil degrading community

**DOI:** 10.1101/595074

**Authors:** Christian M.K. Sieber, Blair G. Paul, Cindy J. Castelle, Ping Hu, Susannah G. Tringe, David L. Valentine, Gary L. Andersen, Jillian F. Banfield

**Affiliations:** Department of Earth and Planetary Science, University of California, Berkeley, CA 94720, USA; Department of Energy Joint Genome Institute, Walnut Creek, CA 94598, USA; Marine Science Institute, University of California, Santa Barbara, CA 93106, USA; Ecology Department, Climate and Ecosystem Sciences Division, Lawrence Berkeley National Laboratory, Berkeley, CA 94720, USA; Department of Biology, St. Mary’s College of California, Moraga, CA 94575, USA; Department of Environmental Science, Policy and Management, University of California, Berkeley, CA 94720, USA

## Abstract

The Candidate Phyla Radiation (CPR) comprises a large monophyletic group of bacterial lineages known almost exclusively based on genomes obtained using cultivation-independent methods. Within the CPR, Gracilibacteria (BD1-5) are particularly poorly understood due to undersampling and the inherent fragmented nature of available genomes. Here, we report the first closed, curated genome of a Gracilibacteria from an enrichment experiment inoculated from the Gulf of Mexico and designed to investigate hydrocarbon degradation. The gracilibacterium rose in abundance after the community switched to dominance by *Colwellia*. Notably, we predict that this gracilibacterium completely lacks glycolysis, the pentose phosphate and Entner-Doudoroff pathways. It appears to acquire pyruvate, acetyl-CoA and oxaloacetate via degradation of externally derived citrate, malate and amino acids and may use compound interconversion and oxidoreductases to generate and recycle reductive power. The initial genome assembly was fragmented in an unusual gene that is hypervariable within a repeat region. Such extreme local variation is rare, but characteristic of genes that confer traits under pressure to diversify within a population. Notably, the four major repeated 9-mer nucleotide sequences all generate a proline-threonine-aspartic acid (PTD) repeat. The genome of an abundant *Colwellia psychrerythraea* population has a large extracellular protein that also contains the repeated PTD motif. Although we do not know the host for the BD1-5 cell, the high relative abundance of the *C. psychrerythraea* population and the shared surface protein repeat may indicate an association between these bacteria.

**Importance:** CPR bacteria are generally predicted to be symbionts due to their extensive biosynthetic deficits. Although monophyletic, they are not monolithic in terms of their lifestyles. The organism described here appears to have evolved an unusual metabolic platform not reliant on glucose or pentose sugars. Its biology appears to be centered around bacterial host-derived compounds and/or cell detritus. Amino acids likely provide building blocks for nucleic acids, peptidoglycan and protein synthesis. We resolved an unusual repeat region that would be invisible without genome curation. The nucleotide sequence is apparently under strong diversifying selection but the amino acid sequence is under stabilizing selection. The amino acid repeat also occurs in a surface protein of a coexisting bacterium, suggesting co-location and possibly interdependence.

## Introduction

Metagenomics data, the DNA sequences from microbial communities, can be used to reconstruct genomes from uncultivated organisms and provide insight into biological processes shaping their ecosystems. The approach has led to the discovery of numerous previously unknown phyla, many of them belonging to the candidate phyla radiation (CPR), which now appears to constitute a major part of the bacterial domain (1, 2). The candidate phylum BD1-5 was first genomically sampled from an acetate-amended aquifer (Rifle, Colorado) (3). The organisms were suggested to have limited metabolism and predicted to be symbionts (possibly episymbionts), but the nature of their associations with other organisms remains a mystery. Wrighton *et al*. (3) predicted that BD1-5 use an alternative genetic code in which the stop codon UGA encodes an amino acid. Following sampling by single cell genomics, BD1-5 were named Gracilibacteria (4). The prediction that UGA codes for glycine in Gracilibacteria was experimentally validated by Hanke *et al.* (5) through proteomic analysis of a sediment enrichment culture. However, the lack of very high quality genomes has limited detailed analysis of the lifestyle of Gracilibacteria and complicated predictions regarding the presence and absence of key metabolic pathways.

Here, we used metagenomic data from a previously performed experiment intended to simulate the Deep Water Horizon (DWH) oil spill (6) to reconstruct the first closed, circular genome (1.34 Mbp) for a Gracilibacteria population. The experiment was inoculated using a water sample collected from the Gulf of Mexico and Gracilibacteria were detected at moderate abundance 64 days after oil droplet addition (Methods). The genome encodes numerous proteins that could not be assigned a potential function, but genes and pathways that are present were easily recognizable. Notably, one hypervariable gene is inferred to encode a protein under strong stabilizing selection, thus likely important for survival. Even for a CPR bacterium, we note an unusual lack of core carbon compound metabolic pathways, including the complete absence of glycolysis and the pentose phosphate pathway. Glycolysis is the major pathway for sugar utilization and is present even in the very small genomes of *Buchnera* and “*Candidatus* Blochmannia”, bacteria that are obligate insect endosymbionts (7), and at least a partial pathway is present in many other symbionts. These observations raise interesting questions regarding how central carbon currencies are acquired and how reducing power is generated and recycled.

## Results and Discussion

### Genome assembly and curation reveals a hypervariable gene

The draft Gracilibacteria (BD1-5) genome binned by Hu *et al*. (6) from sample BD02T64, taken 64 days after the start of the laboratory experiment (Methods), was selected for further curation as it comprised just 6 scaffolds. We verified that these scaffolds cluster tightly together on a tetranucleotide Emergent Self Organizing Map (Figure S1), supporting their derivation from a single genome (8). Protein predictions for all of these six scaffolds required use of an alternative code in which the UGA stop codon is translated as an amino acid. Consistent with prior studies of Gracilibacteria, the genes were predicted using genetic code #25 (UGA translated as glycine (9)). There have been two main ideas proposed to explain how alternative coding arises. The first relates to the low GC content of some (but not all) of the genomes it occurs in. The currently described genome fits this pattern (28.87% G+C). Alternatively, McCutcheon and Moran (10) invoke loss of peptide chain release factor 2 (encoded by prfB), which recognizes UGA codons, to explain the reassignment of stop UGA to tryptophan (code #4) in insect symbionts. Consistent with the hypothesis of McCutcheon and Moran (10), prfB was not detected in the genome of the gracilibacterium studied here, or in any other available BD1-5 genome. However, peptide chain release factor 1 (prfA) was detected, and the gene is widely identified across the CPR.

Prior to read-based curation, the six scaffolds were tentatively condensed into two based on perfect overlaps at scaffold ends. Local assembly errors were removed by curation and unplaced paired reads were used to close gaps. Reads mapped to the scaffolds were visualized in Geneious (11). Notably, a region where two scaffolds were joined based on end overlaps was identified as incorrectly assembled based on the absence of perfect read support. Inaccurate read placements were associated with a hypervariable repeat locus. By manual step-by-step repositioning of paired reads (first placing reads anchored into the repeat locus boundaries, then considering paired read distances and repeat composition) it was possible to generate a representation of the sequence through the locus (Figure 1). Due to the large size of locus compared to the paired read distance, it is impossible to determine the exact number of repeats or if the locus exhibits cell to cell variation in repeat number per locus. However, based on the average sequencing depth, the approximated locus is probably of about the correct length and not highly variable in terms of the number of repeated sequences.

**Figure 1:**
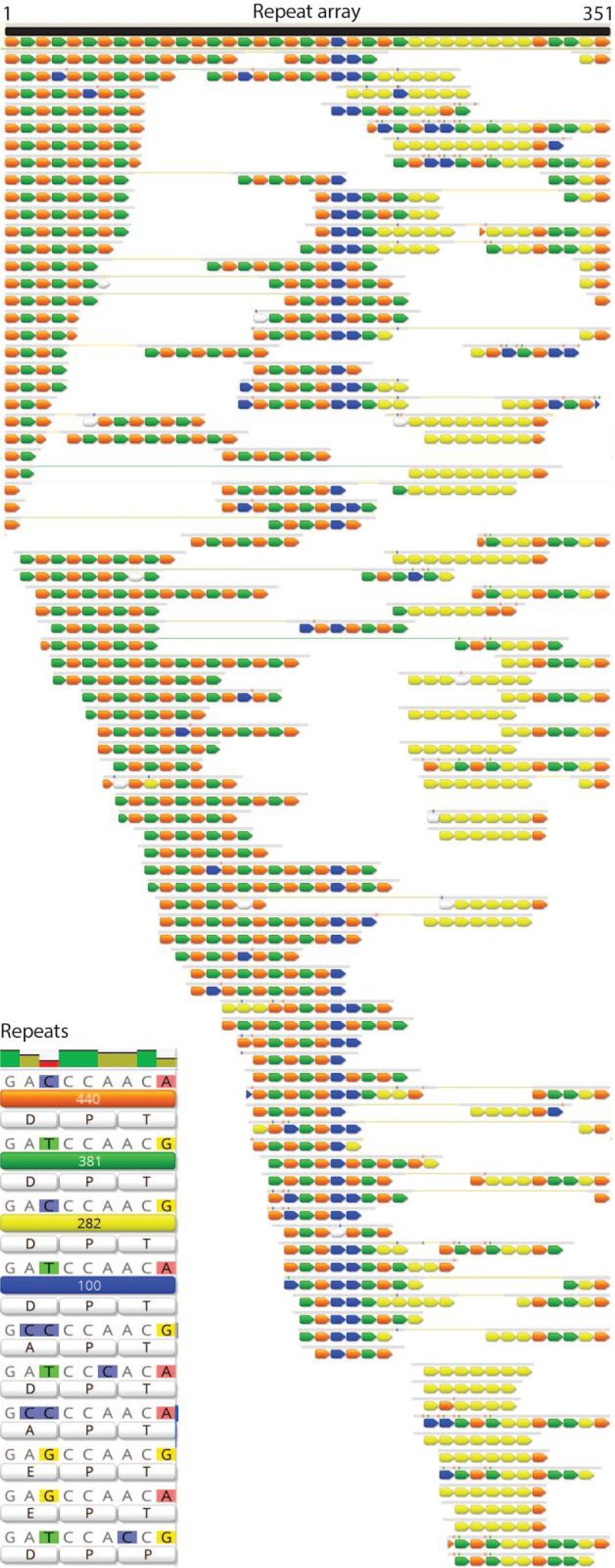
Repeat locus from the BD1-5 genome. Colored arrows represent repeated sequence blocks, the sequences for which are shown in the Repeats insert. Sets of arrows represent reads and reads linked within this region to paired reads are indicated by a thin connecting line.

We verified the final genome path by calculating the cumulative GC skew of the closed chromosome sequence and identified the pattern expected for normal bacterial bidirectional replication (Figure 2). The final assembly comprises 1.34 Mb, 1,243 protein coding genes, 33 tRNA genes and one set of rRNAs (Table 1). According to a RAxML tree based on 16S rRNA genes the closest relative to our organism was sampled from deep sea sediments; other closely related sequences are from marine environments (Figure 3).

**Table 1:** General information about the Gracilibacteria genome.

|  |  |
| --- | --- |
| **Genome statistics** |  |
| Genome size (Mb) | 1.343 |
| GC-content (%) | 28.87 |
| Avg. coverage | 42.26 |
| Protein coding genes | 1243 |
| tRNA genes | 33 |
| rRNA genes | 1 |
| Transcription factors | 29 |
| Secreted proteins (signal peptide) | 66 |
| Small secreted proteins (< 300 aa) | 24 |
| Non-classically secreted proteins (no signal peptide) | 104 |
| Small Non-classically secreted proteins (< 300 aa) | 61 |
| Transmembrane proteins (>3 TM domains) | 122 |
| Transporters | 481 |
| ABC transporter related proteins | 11 |
| Amino acid permeases | 34 |

**Figure 2:**
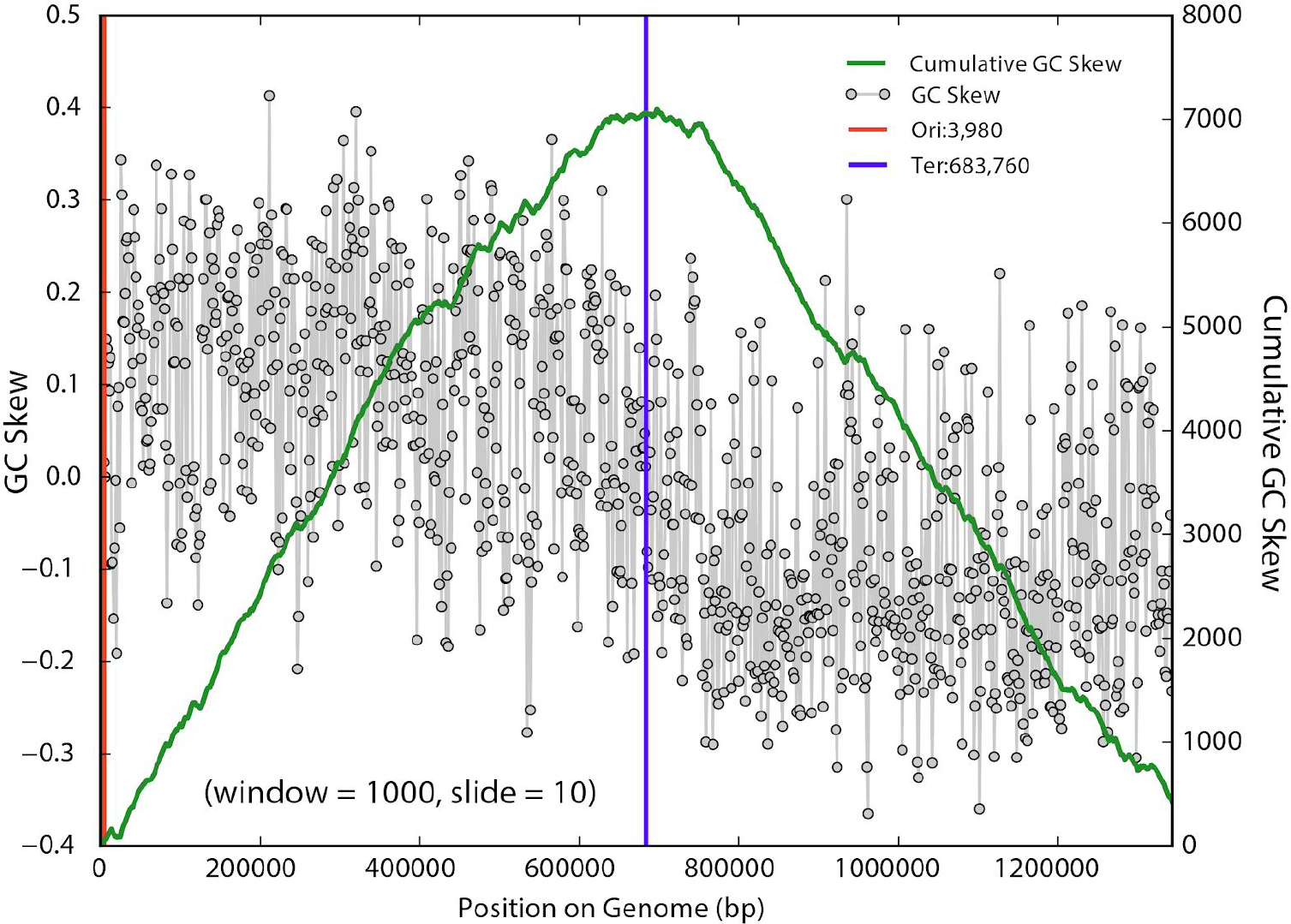
Diagram showing the GC skew (grey dots) and calculated cumulative GC skew (green line) across the finished BD1-5 genome. The pattern is typically of a correctly assembled genome of a bacterium that undergoes bidirectional replication from origin to terminus.

**Figure 3:**
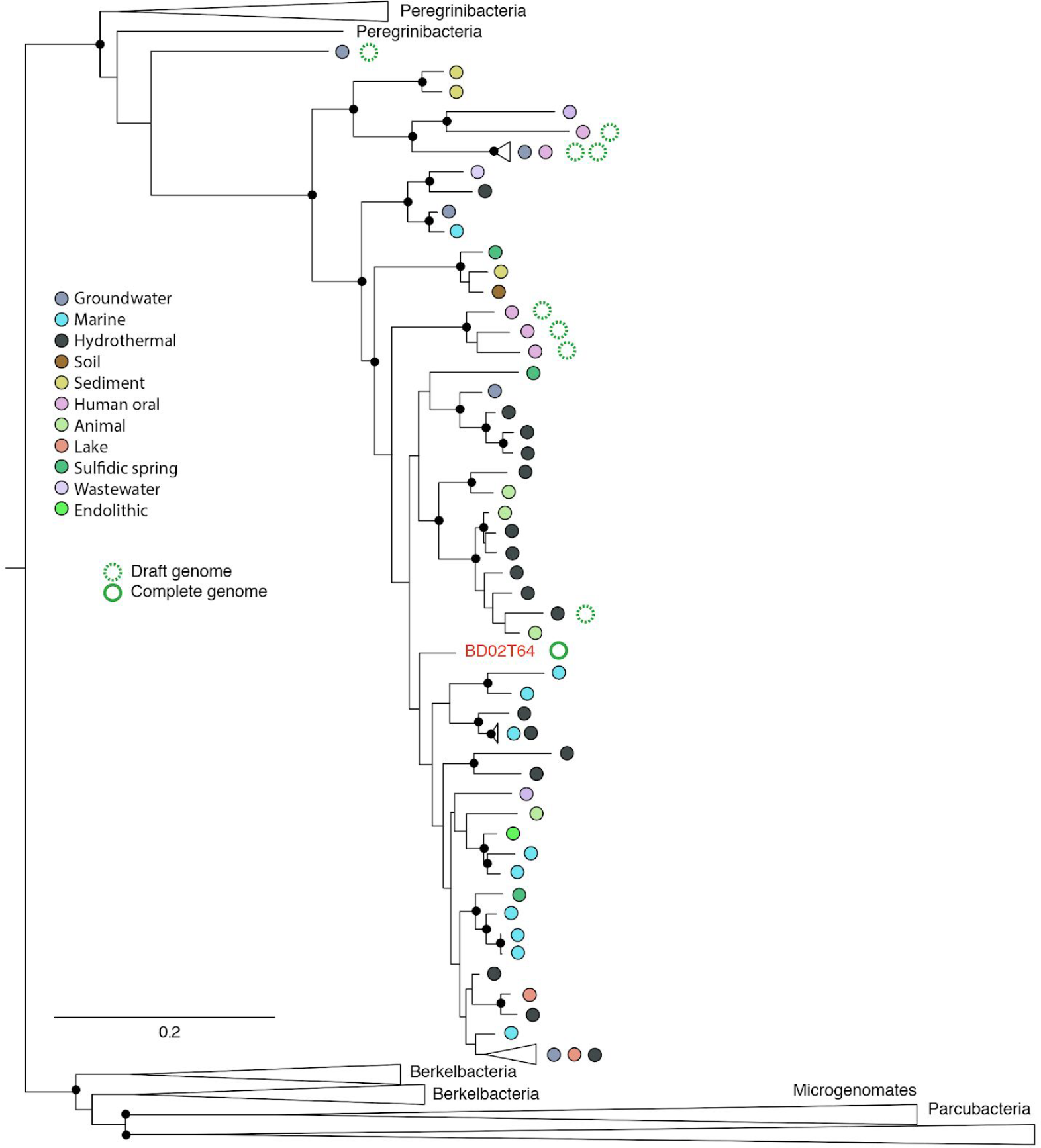
Phylogenetic placement of the Gracilibacteria genome from sample BD02T64 reported here. The 16S rRNA tree was constructed using the maximum likelihood method RAxML. The small black circles indicate nodes with Maybe just list support values > 70% bootstrap support. 16S rRNA genes retrieved from genomes are indicated by green circles. Dotted circles represent published draft genomes and the full circle indicates the finished and curated genome from this study. Colored circles indicate the type of ecosystem from which sequences were obtained. The full tree file is provided in the Supplementary Online Materials.

The predicted amino acid sequence of the BD1-5 gene containing the repeat region is shown in **Display Item Supplementary 1A**. Some repeat types occur in blocks and some repeat types alternate, but overall the most striking feature of the locus is the high level of cell to cell heterogeneity (Figure 1). Variant calling in reads mapped to the full length protein identified 17 synonymous single nucleotide substitutions and zero non-synonymous substitutions (**Table S1**), with the exception of instances occurring only on a single read. In fact, this gene contains the highest proportion of synonymous substitutions in any in the BD1-5 genome.

Within the repeat region, the incidence of synonymous vs. non-synonymous substitutions is shown in Figure 1 (insert). The four main repeat nucleotide sequence variants are indicated in orange, green, yellow and blue, along with their translated sequences. Single incidence sequences are indicated by white bars. Notably, the nucleotide sequences of the four major repeat variations all translate to the tripeptide amino acid motif: PTD. Given that the Gracilibacteria population cells share near-identical nucleotide sequences genome wide, except within this specific locus, we infer that the repeat bearing protein may be under strong pressure to evolve at the nucleotide level. If cells acquire non-synonymous substitutions in the repeat protein, they are apparently strongly selected against.

The PTD repeat motif is found in hypothetical proteins and predicted surface proteins of a few other organisms, including some eukaryotic sporozoite surface protein 2-like. A secondary structure prediction of the BD1-5 protein suggests only beta sheet and coils, with the repeat motif in a coil region. We predict a single N-terminal transmembrane domain and extracellular localization of the remainder of the protein sequence, including the repeat region (Figure S2).

We investigated codon usage in the repeat gene. The codons for D are GAC and GAT, with usage of 6.2:6.7 in the repeat gene, whereas the expected genome-wide incidence is GAC: GAT = 0.83:4.79. Synonymous substitutions within the repeats could cause ribosome pausing and modulate rates of protein folding (12, 13). Thus, the atypical codon use in this region might indicate selection for frequent translational pausing. The codons for P are coded for by CCA, CCG, CCT, and CCC, with an expected incidence of 1.3:0.13:0.89:0.13, whereas the repeat gene has an incidence of 11.82:0.25:1.23:0. Thus, there is evidence for strong selection for the CCA codon (considered further below). In the third position of the tripeptide, T can be ACT, ACC, ACA, ACG with an expected incidence: 2.1:0.41:2.4:0.36. However, within the repeat gene ACT, ACC, ACA, ACG occur with an incidence of 4.19:0:6.9:6.4. The reliance on rare codons may again indicate selection for frequent translation pausing.

If it is advantageous for coexisting cells to have highly variable rates of translation, one might expect that the sequences would make maximal use of the available codons. Counter to this, we see reduced codon diversity. Thus, we considered that variation in the secondary structure of the RNA sequence in the repeat array may be selected for. In the secondary structure prediction for the repeat region we note the periodic alternation of stems, comprising mostly Watson-Crick base pairs, and loops (Figure S3). Notably, the CCA codon (specifically the first C) is at the base of the bubbles and closes them, paired to Gs from either the first base of the first codon or the last base of the third codon. Stem loops impact RNA folding, can stabilize mRNA and provide recognition sites for RNA binding proteins. We speculate that nucleotide variation may impact the translation rate of this gene and lead to variation in the fitness of different population members.

We searched the genomic region flanking this gene but did not identify a known mechanism for site-directed mutagenesis within the repeat locus. Genes with functions linked to DNA repair and recombination are found in close downstream proximity (uvrC excinuclease (5798 bp downstream), an exodeoxyribonuclease III (8336 bp downstream), and DNA recombination-mediator protein, DprA (30798 bp downstream). Perhaps this organism possesses a DNA mutator, which mediates targeted diversification in the repeat locus (e.g., UmuCD). It is unlikely that the organism is deficient in repair enzymes, as sequence variation is not elevated elsewhere in the genome. Perhaps nucleotide heterogeneity arose due to suppressed proofreading in this region, but we have no explanation for how this might have occurred.

In the current study, it is difficult to evaluate locus length variation because read lengths are short compared to the length of the repeat arrays. Locus length variation is expected, given the presence of perfect repeat arrays. In bacterial genomes, repeat regions may expand and contract due to either recombination or slipped-strand mispairing (SSM; (14, 15)), resulting in population variability in terms of tripeptide motifs that may impact three dimensional protein structure and ligand binding. The relationship between microsatellite length and point mutation has been described elsewhere and generally predicts that as a locus expands, base substitutions accumulate and suppress further SSM (16, 17). If SSM is undesirable, it is advantageous to include nucleotide variants that offset repeat pairing and thus prevent slippage.

Examination of the 5’- and 3’-untranslated regions flanking the repeat gene uncovered two sequences capable of forming stem-loops with notably long stems (14-15 bp) and 4-6 bp loops (Figure S4). As DNA or RNA structures, these stem-loops may play a role in recombination or as transcriptional regulation signals for the repeat-containing gene, respectively.

Some reads mapped to the BD1-5 repeat region had paired reads that were not placed in that genome. Comparison of the non-repeat regions of these reads and the sequences of their unplaced paired reads to the genomes of other community members revealed 100% nucleotide matches to a region on BD02T64_scaffold_179, part of a draft *Colwellia psychrerythraea* genome (BD02T64_Colwellia_psychrerythraea_38_180_partial). Thus, we concluded that a region within the genome of this abundant population has the same PTD repeat as found in the Gracilibacteria protein (**Display Item Supplementary 1B**). After curation of the region, the *C. psychrerythraea* protein is predicted to be 3459-amino acid in length with a signal peptide and extracellular localization, possible galactose/carbohydrate-binding domains, pectin lyase/virulence domains and parallel beta helix repeats. The repeat occurs within a structure that otherwise consists of a mixture of alpha helices and beta sheets but is in neither of these.

Within the *Colwellia* protein, reads carry up to 11 repeats (Figure S5). As for BD1-5, it is impossible to detect variation in repeat number in each cell due to the read length limitation, but one read has only five repeats. In virtually all cases the nucleotide repeat is encoded by a single 9-mer (yellow), This 9-mer is prominent towards the end of the Gracilibacteria repeat region. The loci in both genomes terminate with the same 9-mer (orange in Figure S5). The essentially perfect repeated sequence would make this region prone to replication slippage, leading to cell-to-cell variation in the number of tripeptides in the protein.

Interestingly, several of the Gracilibacteria proteins encoded immediately adjacent to the variable PTD protein have the highest similarity to proteins in organisms that are not part of the CPR. One is most similar to a protein from *Colwellia psychrerythraea*, although the percent amino acid identity is low (~53%). To rule out chimeric assembly of sequence from another bacterium in this genomic region we confirmed the expected alternative coding throughout (and paired read placements were verified during the main curation phase). Thus, the region encoding the Gracilibacteria variable repeat gene and adjacent genes may have been acquired from a bacterium related to *Colwellia psychrerythraea*.

### Metabolic analysis

The biosynthetic pathways easily recognizable in the genome are for ribosome-based protein synthesis, nucleic acid synthesis and interconversion, DNA repair, peptidoglycan production, secretion, pilus production and cell division. However, as for other CPR, this gracilibacterium appears to lack the ability to synthesize lipids needed for construction of the cytoplasmic membrane (and, there is no pathway for synthesis of Lipid A required for a Gram-negative cell envelope). Thus, these cells are predicted to be either symbionts or closely dependent on other community members for key building blocks. The genome lacks a CRISPR-Cas system for phage defense, but has a restriction modification system that may serve this purpose (18, 19). Absent are almost all pathways for amino acid synthesis, leading us to conclude that amino acids needed for protein biosynthesis are derived through breakdown of externally-derived peptides. Many different types of peptidases and proteases are available for this process.

For nucleic acid synthesis the genome encodes the steps required to interconvert nucleotides. We also identified most of the genes required for biosynthesis of purines and pyrimidines from glutamine and aspartate; these genes are relatively uncommon in CPR. Inosine monophosphate (IMP) can be converted to adenosine monophosphate (AMP), adenosine diphosphate (ADP), adenosine triphosphate (ATP) and incorporated into RNA and DNA. Enzymes were also identified to interconvert forms of GDP and GTP. Genes of the one carbon pool by folate pathway were identified, enabling transfer of C1 groups during nucleotide metabolism, but genes for folate biosynthesis were not identified.

Perhaps the most surprising feature of this bacterium is the complete lack of genes for glycolysis and the pentose phosphate pathway. At least partial pathways are present in other Gracilibacteria, and the first reported genomes have full pathways to convert glucose to pyruvate and fermentation-based metabolisms (3). More broadly, at least parts of these pathways are present in the most minimal CPR genomes. However, this is the first genome from a major subgroup within Gracilibacteria (Figure 3), so it remains to be seen whether this is a common trait. The absence of these pathways raises two questions (1) the nature of central carbon metabolism in these organisms, and (2) how ATP, NADH, NADPH and ferredoxin are reduced and recycled.

Potentially addressing the first question, we identified a variety of pathways for production of central carbon currencies. We identified a putative two subunit ATP citrate (pro-S)-lyase [EC:2.3.3.8] (genes 1051, 1052), a complex rarely detected in CPR. This annotation (vs. citrate synthase) was supported by HMM homology and the presence of the active site residues GHAGA (20). Via this complex, citrate can be converted to acetyl-CoA and oxaloacetate. Citrate may be obtained from external sources via two putative citrate transporters. Intriguingly, both ATP citrate (pro-S)-lyase subunits are most similar to predicted proteins in archaea, suggesting their acquisition via lateral gene transfer. We predict that oxaloacetate derived from breakdown of citrate is converted to pyruvate via a 2-oxoacid ferredoxin oxidoreductase (OFOR). Pyruvate can also be produced from phosphoenolpyruvate via a pyruvate kinase, and from malate [1.1.1.38] and serine [4.3.1.19]. Overall, amino acids scavenged from the environment appear to feature prominently in the metabolism of this gracilibacterium, and some are converted into the nitrogen and carbon storage compound cyanophycin.

Addressing the second question, we identified many reactions that oxidize or reduce energy currencies via transformation of small carbon compounds. Specifically, pyruvate conversion to acetyl-CoA via OFOR consumes NADH while reducing ferredoxin. The ferredoxin may be reoxidized via either a cytoplasmic or membrane bound ferredoxin reductase (FNR) that also converts NADP+ to NADPH. NADH may be regenerated in the production of pyruvate from serine or malate. Like citrate, malate may be obtained from external sources. Other reactions, such as those involved in peptidoglycan synthesis and interconversion of tetrahydrofolate compounds, also interconvert energy currencies. Enzymes that respond to oxidative stress response also may provide electron sinks.

Many CPR generate ATP via substrate level phosphorylation reactions that produce compounds such as acetate, but genes for production of these short chain fatty acids were not identified. ATP required for DNA and RNA biosynthesis may be formed via the F-ATP synthase complex (Complex V). Given the lack of an electron transport chain that could pump protons, proton motive force could be stolen from attached host cells if tight junctions are formed (19, 21). Such close physical associations have been reported for another CPR bacterial group, Saccharibacteria (TM7), which attach to host Actinobacteria cell surfaces (22). Alternatively, proton motive force could be generated by cytoplasmic draw down of H^+^ via reactions involved in breakdown of amino acids and other compounds, Na^+^/H^+^ antiport, or consumption of H^+^ by superoxide dismutase. The ATP synthase may also be used reversibly to generate proton motive force (as suggested by Wrighton *et al*. (3)), but no complexes were identified that could make use of the generated PMF. Specifically, there is no indication of hydrogenases, which occur in some other CPR. Lacking also are other electron transport chain components such as NADH dehydrogenase, succinate dehydrogenase and cytochrome-c reductase/oxidase and most steps of the tricarboxylic acid cycle (Figure 4).

**Figure 4:**
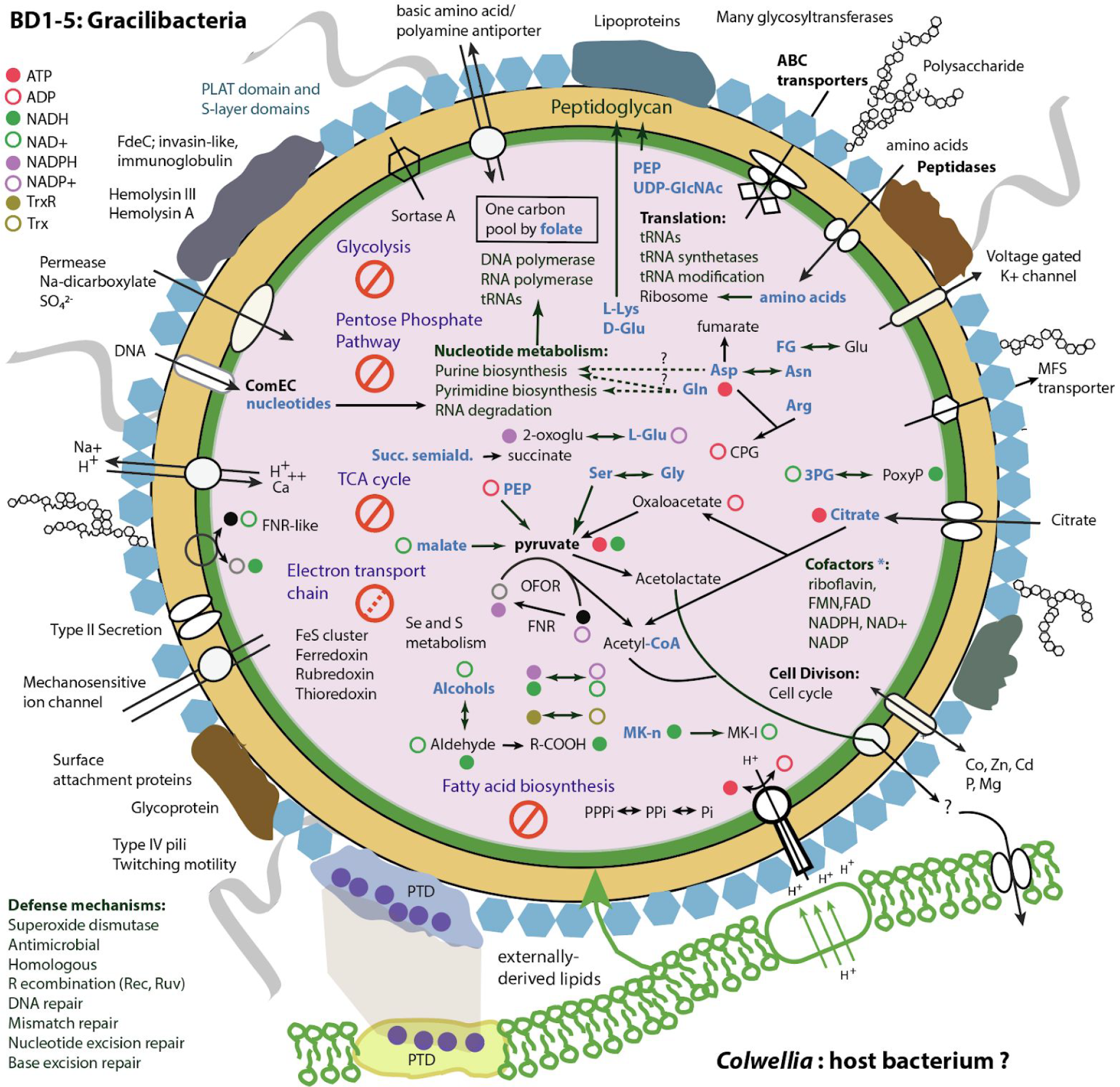
Cell cartoon depicting a reconstruction of the metabolism of the gracilibacterium. Bold text indicates prominent functions, blue text indicates resources inferred to be externally derived. The * indicates that reactions for biosynthesis of cofactors require a precursor compound. Abbreviations: PEP: phosphoenolpyruvate, UDP-GlcNAc: UDP-N-acetyl-alpha-D-glucosamine, OFOR: 2-oxoacid ferredoxin oxidoreductase, 3PG: 3-Phospho-D-glycerate, 3-PoxyP: 3-Phosphonooxypyruvate, 2-oxoglu: 2-Oxoglutarate, FNR: ferredoxin reductase, PPPi, PPi, Pi: phosphate compounds interconverted by inorganic pyrophosphatase, Mk-n: metaquinone, Mk-l: metaquinol, Succ. semiald.: succinate semialdehyde, L-Glu: L-glutamate, R-COOH: a carboxylic acid, CPG: cyanophycin. FG: N-formyl-L-glutamate. PTD is a tripeptide repeat.

A variety of transporter types were predicted, presumably addressing the need to acquire compounds from other cells or detritus. There are many hypothetical membrane-associated proteins with multiple transmembrane (TM) domains that also may serve a transport role. Overall, 122 transmembrane proteins (>3 transmembrane domains) and 481 transporter proteins were identified (Table 1). The genome encodes an intriguing 990 aa protein predicted to contain 32 transmembrane domains (Gene 860). A large scale analysis of TM rich proteins in NCBI nr revealed that very few have 32 or more TM domains, and only a few related proteins are known (mostly in other Gracilibacteria). The function of this enigmatic protein is uncertain as the only domain predicted is DUF2339 (hypothetical membrane protein).

A notable feature of the Gracilibacteria genome is the prominence of secretion mechanisms and secreted proteins. We identified 66 such proteins using a combination of three methods to predict signal peptide mediated export, of which only 24 are shorter than 300 amino acids. A further 104 proteins are predicted to be secreted via non-classical pathways that do not use a signal peptide. Of these, 43 are larger than 300 amino acids. In addition to a sortase (typically found in Gram-positive bacteria and common in CPR), we identified genes of the type II and IV secretion pathways that are generally associated with Gram-negative bacteria, including multiple copies of SecA,D,F,Y,E,G). SecYEG form the central translocase across the inner membrane, SecA guides proteins to the translocase channel and is the ATPase and SecF promotes release of the mature peptide into the periplasm. Thus, the identified components provide the functions required for secretion in non-Gram-negative bacteria. Intriguingly, 15 general secretion protein G proteins (gspG, alternatively pulG) are predicted, as well as gspE. These are large proteins, on average 528 aa in length. GspG is the major pseudopilin present in a pseudopilus and GspE is an ATPase involved the assembly of the pseudopili. In addition, we identified around 12 type IV pilus assembly protein subunits, some in multi-copy. Type IV pili allow the transfer of genetic material PilV,C,B,W and are involved in twitching motility (the genome also has two pilT). pilD (leader peptidase) was also identified. We did not identify PilQ, consistent with lack of outer membrane. Pili may be involved in attachment and inter-organism interactions, as well as uptake of DNA. Competence genes were also identified (19, 21, 23).

From the perspective of the cell envelope, the biosynthesis pathway for peptidoglycan is complete, although the requirement for precursor UDP-N-acetylglucosamine from external sources is predicted. Predicted are genes to convert phosphorylated isoprenoid into a precursor for peptidoglycan, but the genome lacks the archaeal mevalonate and bacterial MEP (2-*C*-methyl-D-erythritol 4-phosphate) pathways. It has geranylgeranyl diphosphate synthase, but the reason is unclear. In addition, we identified three genes that degrade L-lysine and D-glutamate that may feed intermediates into two different steps within the peptidoglycan biosynthesis pathway. The genome encodes many genes for polysaccharide synthesis (e.g., #444-460) and for proteins with S-layer domains. Thus, we anticipate a cell wall containing peptidoglycan with a periodic surface layer, many and potentially diverse pili and a variety of large extracellular proteins and polymeric substances (Figure 4). Interestingly, some S-layer proteins may have toxin domains (e.g., 1226, predicted to have polycystin-1, lipoxygenase, alpha-toxin domains). Other large proteins have annotations suggestive of hostile interactions with other organisms (e.g., insecticidal toxin complex protein TccC) and there is a predicted invasin domain in one large protein in the genome.

In terms of the ability to respond to environmental conditions, the genome encodes at least four relA/spoT domain proteins, three of them encoded sequentially and one larger multi-domain protein encoded elsewhere. These may function in response to nutrient limitation. Also identified are two 8-oxo-dGTP diphosphatase genes to prevent misincorporation of the oxidized purine nucleoside triphosphates into DNA and proteins with antioxidant functions, including superoxide reductase and enzymes to reduce oxidized methionine.

We conclude that the inferred putative symbiotic lifestyle of Gracilibacteria differs in notable ways from those of other obligate host-associated organisms. The genome size is large, compared to those of most obligate host-associated organisms (usually <1 Mbp in length; (24)). Host-associated bacteria that have experienced moderate genome reduction retain genes for synthesis of fatty acids and peptidoglycan (but not for LPS or phospholipids) whereas those that have undergone extreme genome reduction have essentially no genes for cell envelope biosynthesis (10). In contrast, the gracilibacterium seems to rely entirely on externally derived fatty acids. It retains genes for regulation of gene expression (e.g., two-component systems and various transcriptional regulators), DNA repair and homologous recombination, whereas these genes are often lost in symbionts (7). Overall, the genomic features of this gracilibacterium only overlap partially with those of host-associated bacteria, which have experienced rapid genome decay.

## Conclusions

Among the most intriguing aspects of the Gracilibacteria genome studied here is the variable nucleotide locus that encodes a conserved tandem PTD tripeptide repeat protein. The gene appears to be under selective pressure to preserve this sequence, as nucleotide variation is localized to this repeat locus almost exclusively as synonymous codons. We infer that the protein has a function strongly tied to the fitness of this organism. The PTD repeat sequence also occurs in coexisting *Colwellia* that became abundant late in the experiment (6) when the gracilibacterium was detected. It is unlikely that co-occurrence of the repeat is a coincidence, as this sequence is relatively uncommon, even in public databases. Given this, and its likely function as an extracellular protein potentially involved in attachment, we speculate that the selective pressure may relate to either host attachment or host-symbiont co-localization (Figure 4). The observations motivate enrichment experiments targeting *Colwellia* to determine if co-cultivation with Gracilibacteria can be achieved.

The gracilibacterium studied here is also fascinating in terms of its unusual metabolic platform. Based on its predicted gene inventory, it is inferred to adopt the lifestyle of a scavenger or symbiont of some type (possibly as a parasite). Certainly, it requires an external source of building blocks, including lipids, amino acids, citrate and malate. In the enrichment experiment designed to simulate the Deepwater Horizon oil spill, glucose-based compounds are not expected to be in high abundance, nor are amino acids. There is no indication that the gracilibacterium can metabolize complex oil-derived compounds. Thus, we predict that the relevant resources are probably bacterial compounds released by cell lysis (e.g,. amino acids, small organic molecules, lipids and cofactors or cofactor precursors) and those that leak from cells of coexisting oil-degrading bacteria (e.g., alcohols and aldehydes). These resources may be processed by this gracilibacterium and the byproducts excreted, provisioning the associated organisms with compounds such as acetyl-CoA, fumarate, succinate or acetolactate. Based on its inferred lifestyle and its phylogenetic placement within a major distinct clade (Figure 3), we propose the name Gracilibacteria (phylum), Gracilibacter (class), *Detritibacteriales* (order), *Detritibacteriaceae* (family), *Detritibacteria* (genus) *gulfii* (species), reflecting its likely dependence on detritus and enrichment in a sample simulating the Gulf oil spill.

## Supporting information

16S rRNA RAxML tree in Nexus format

## Data Availability

The genome, with functional annotation, can be accessed: https://ggkbase.berkeley.edu/BD02T64/organisms/60439

The genome sequence has been deposited in NCBI under accession TBD (Submission ID: SUB5359196).

## Methods

### Genome assembly and annotation

The original study of Hu *et al*. (6) involved seawater samples collected from a depth of 1100-1200 m in the Gulf of Mexico in 2014. The sample derived from a region impacted by the Deep Horizon oil spill in 2010, but there were no oil spill-derived hydrocarbons detected at the time of sampling. However, hydrocarbons seeps occur naturally in the general area. The in situ cell density was estimated at ~ 5.0e+5 cells/ml. A volume of 630 L was returned to the surface and amended with unweathered Macondo oil (MASS oil 072610-03) at a concentration of 0.2 ppm to sustain microbial activity, maintained in the dark at 5 °C while the sample was transported to the laboratory. In the experiment described previously, samples were incubated for up to 64 days in 2-L bottles at 4° in the dark at 0.75 rpm on a rotation carousel system. Macondo crude oil was added to the seawater in 10-um droplets to a final concentration of 2 ppm and 0.02 ppm Corexit EC9500A dispersant (Nalco). Replicate oil-amended bottles were destructively sampled at 6, 18, and 64 d of incubation for metagenomics.

The methods for the metagenomic assembly of the genome of the BD1-5 described here, as well as the draft *Colwellia* genome, are reported by Hu *et al*. (6). In the current study, genome curation was conducted in Geneious (11). Curation involved visualization and validation of paired read placements throughout. Local assembly errors were identified as regions lacking perfect read support. Gaps were inserted in these regions and unplaced paired reads used to fill the gaps. In repeat regions, some reads were improperly placed and paired reads were missing. Curation of these regions was similar to that for local assembly errors, except reads had to be relocated manually to achieve the most parsimonious path. The same approach was used to curate the *Colwellia* genomic region that shared the same repeat sequences. After completion, the assembly was checked for repeats longer than the paired read distance using and GC skew and cumulative GC skew calculated using previously published methods (25).

Genes of the curated, circularized BD1-5 genome were re-predicted using Prodigal (26) with genetic code 25 (-g 25). Functional annotations were done using the ggKbase annotation pipeline (http://ggkbase.berkeley.edu), which searches homologs of predicted genes in the databases of Kegg (27), UniRef (28) and UniProt (29) using USEARCH (30). Amino acid sequences of genes without a significant hit were further annotated using HHblits (31) and the uniprot20 (29) database. In addition, individual genes were interrogated using HHMer (32), HHpred (33), Interproscan (34), Swiss Model (35) and blastp domain analysis. We predicted secreted proteins using psortB (36), signalP (37), and PrediSi (38) with gram negative and gram positive prediction model, respectively. From the six predictions we selected proteins that were identified as secreted proteins by at least three different predictions (coming from at least two independent methods). We applied SecretomeP (39) to predict non-classically secreted proteins without signal peptide. Additionally, we removed proteins with more than one transmembrane domain predicted by TMHMM (40). Transporter proteins were predicted using TrSSP (41).

RNA secondary structure within the repeat locus was determined using YASPIN (42), and DNA secondary structure was predicted using MFold (43) for putative stem-loops flanking the BD1-5 repeat gene. Tertiary structure prediction of the BD1-5 repeat protein was performed using I-TASSER (44).

### Phylogenetic tree

16S rRNA gene sequences were aligned using SSU-align (45), trimmed manually. We calculated the phylogenetic tree using the maximum likelihood algorithm RAxML (46) on the CIPRES (47) web server in choosing the GTRGAMMA model and autoMRE to automatically determine the number of bootstraps.

### Nucleotide Variation and Codon Usage Analysis

We determined single nucleotide variants using VarScan (48), with the following parameters: c10, q30, fr0.05. This set of nucleotide variants were then assessed to determine non-synonymous versus synonymous substitutions within each coding region of the BD1-5 genome. For each gene, we determined the number of codon positions corresponding to an amino acid substitution based on genetic code #25 (Gracilibacteria), versus those resulting in no amino acid change, counted as either non-synonymous or synonymous, respectively. A codon usage profile was generated in Python (v.2.7.3) using the Biopython SeqUtils package. Synonymous codon usage was assessed in the repeat-rich gene for comparison with the average codon usage of all genes in the Gracilibacteria genome.

## Acknowledgements

Support for this research was provided by the Chan Zuckerberg Biohub (to JFB), the Emerging Technologies Opportunity Program of the US Department of Energy (DOE) Joint Genome Institute, a DOE Office of Science User Facility, supported under contract no. DE-AC02-05CH11231. Support was provided by DOE grant no. DOE-SC10010566 and National Institutes of Health grant no. 5R01AI092531. Also, the Center for Dark Energy Biosphere Investigations supported this work (C-DEBI). We thank Brian Thomas for bioinformatics support and Spencer Diamond and David Low for helpful suggestions.

## Author contributions

The previously reported cultivation experiment was designed by GA and PH, and genome assembly and binning were done by PH and CS. The BD1-5 manual genome curation and repeat locus resolution was performed by JFB. Genome wide inventory and protein localization analyses were done by CS. Metabolic analyses were conducted by JFB, CC and CS, with input from BP and DV. Phylogenetic analysis was carried out by CS. Codon and repeat region analyses were performed by BP. Secondary structure analyses were done by BP and JFB. JFB and CC wrote the paper, with input from CS, BP and DV. All authors read and commented on the paper.

## Supplementary Data and Figures

**Figure S1:**
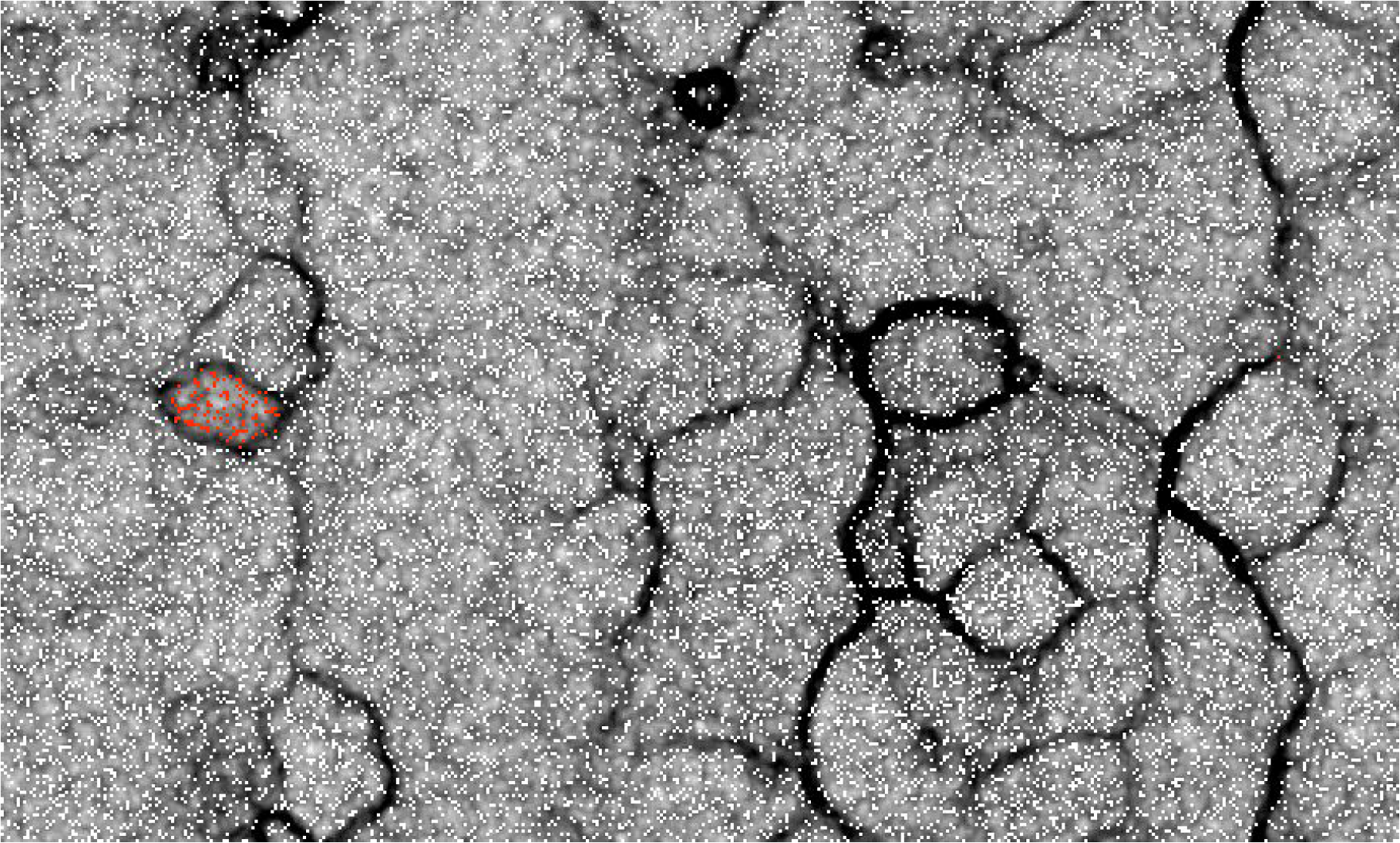
Emergent self organizing map of showing clear clustering together of genome segments (red dots) of the scaffolds assigned to the BD1-5 bin from sample BD02T64 based on tetranucleotide frequencies.

**Display Item Supplementary 1A:**
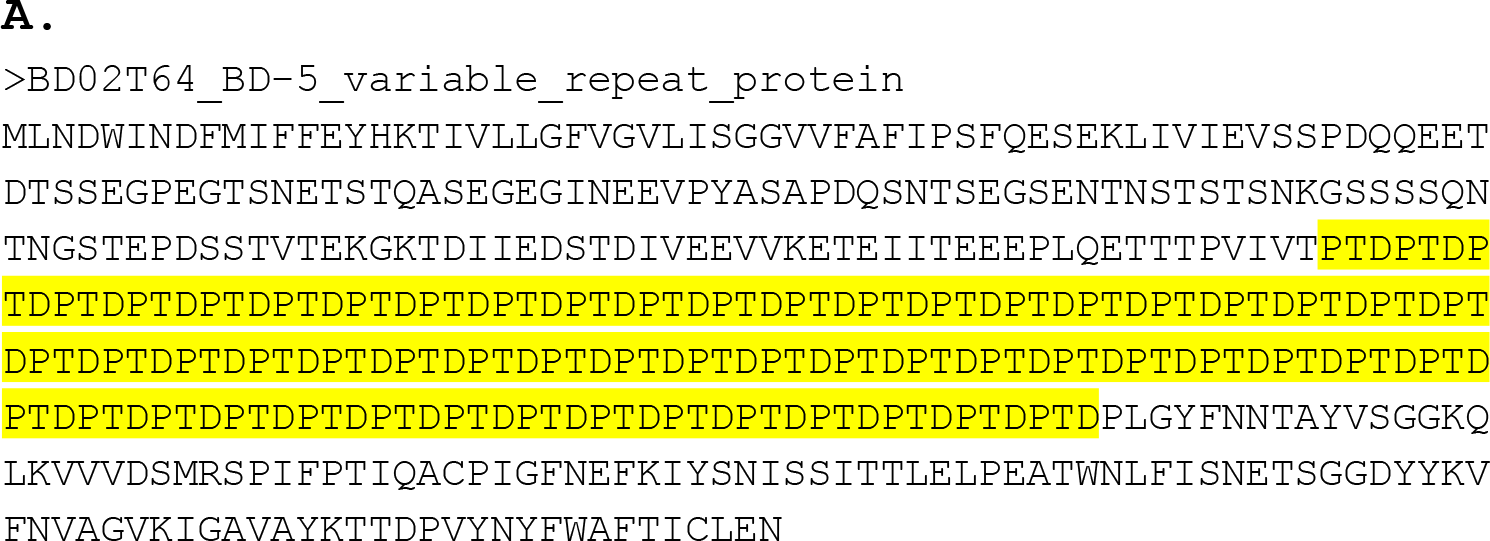
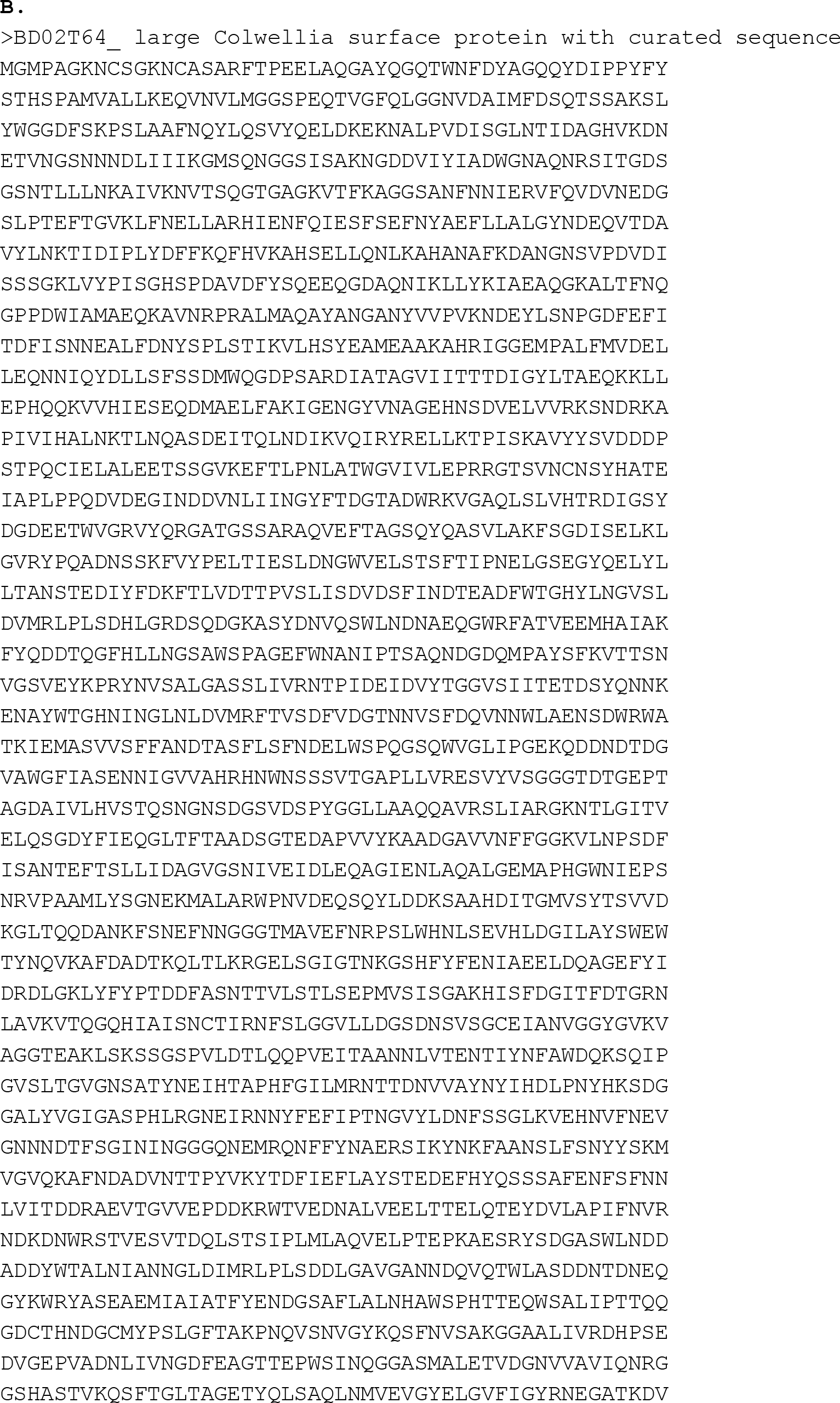
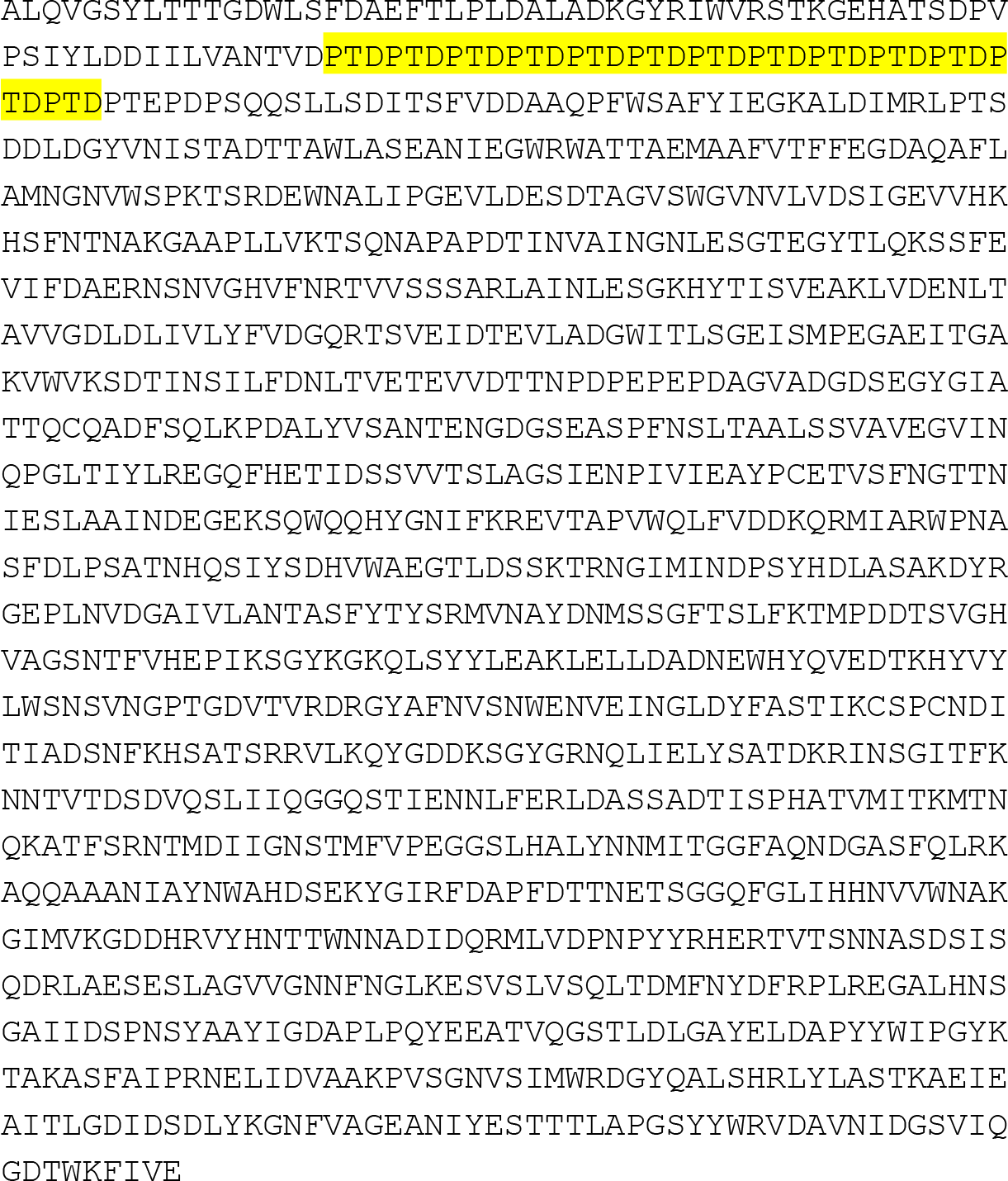
Sequence of the Gracilibacteria repeat protein. The sequence is approximate in terms or repeat number and read arrangement due to the limitation of the read length (see Figure 2). Interestingly, Candidatus Gracilibacteria bacterium HOT-871 (RefSeq: GCF_002761215.1) does not have this protein. **B**. Curated 3,459 aa sequence of the large *Colwellia* surface protein that also contains the PDT repeat. The sequence is predicted to have two carbohydrate binding domains and a central right handed beta helix region. The repeat region appears just after the second carbohydrate binding domain.

**Figure S2:**
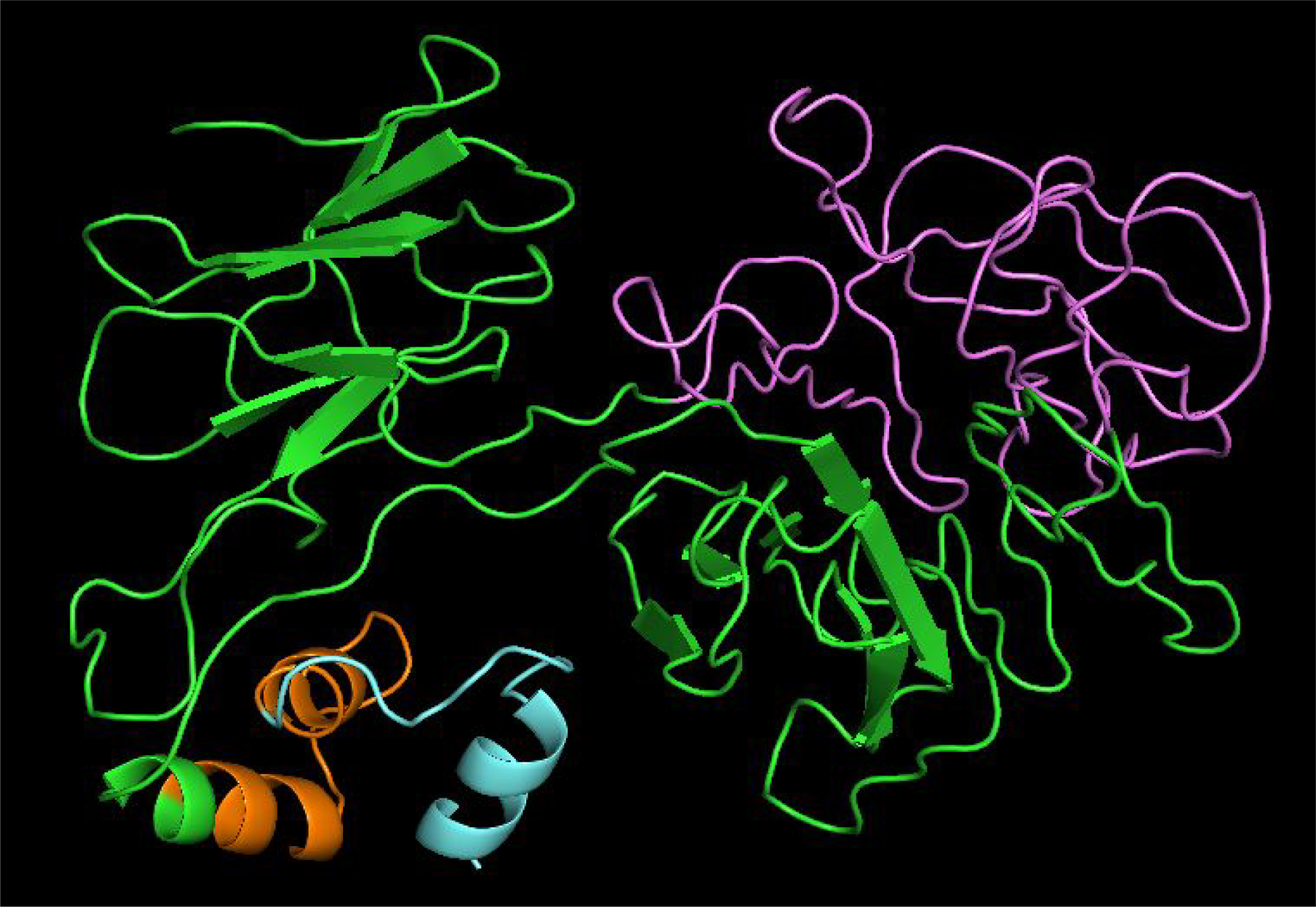
Tertiary structure prediction of BD1-5 repeat protein by I-TASSER showing variable repeat region in purple, transmembrane region in orange and non-cytoplasmic region in turquoise.

**Figure S3:**
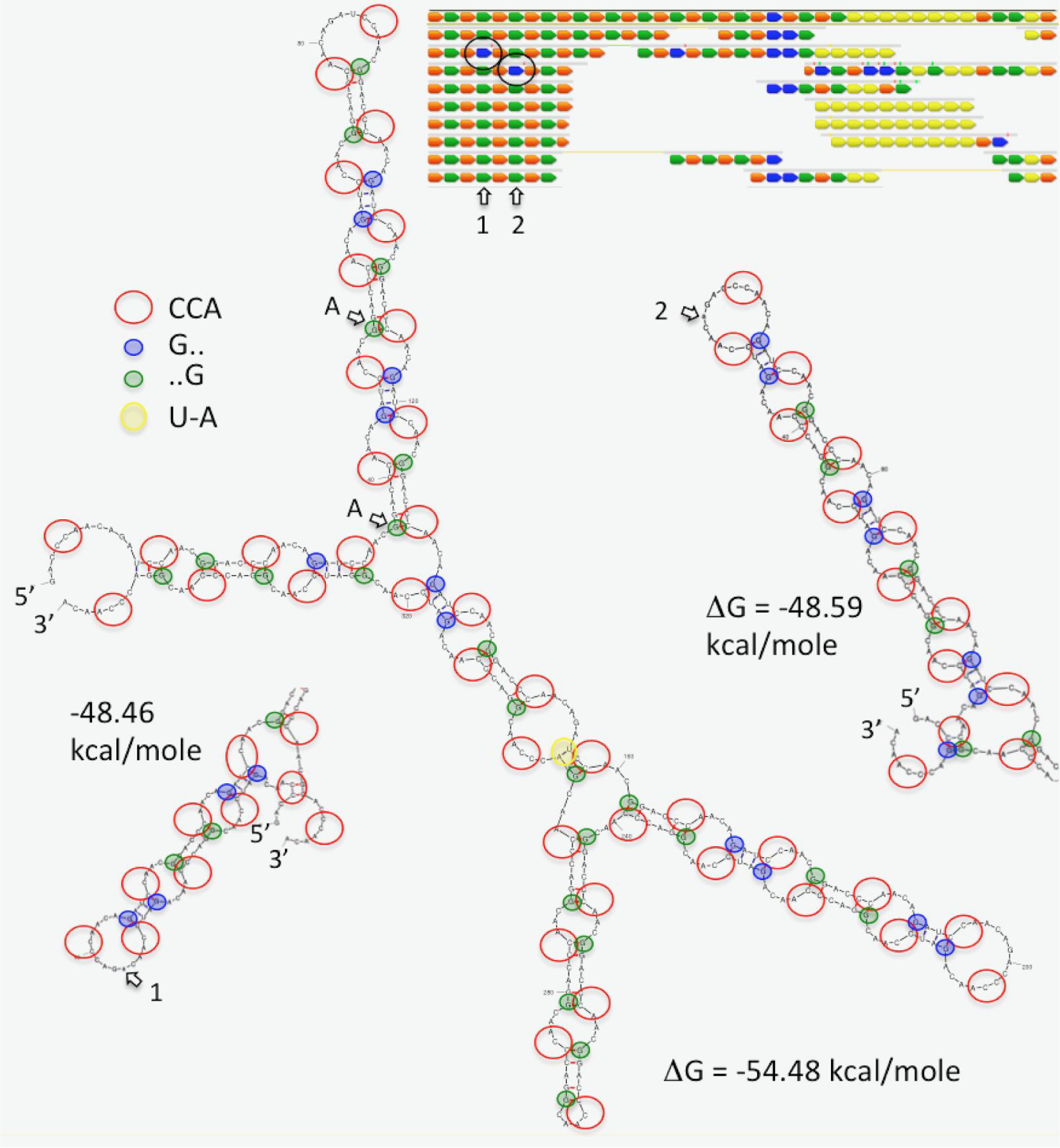
Secondary structure of repeat locus. The consequence of substitutions 1 and 2 on the secondary structure are indicated in local regions.

**Figure S4:**
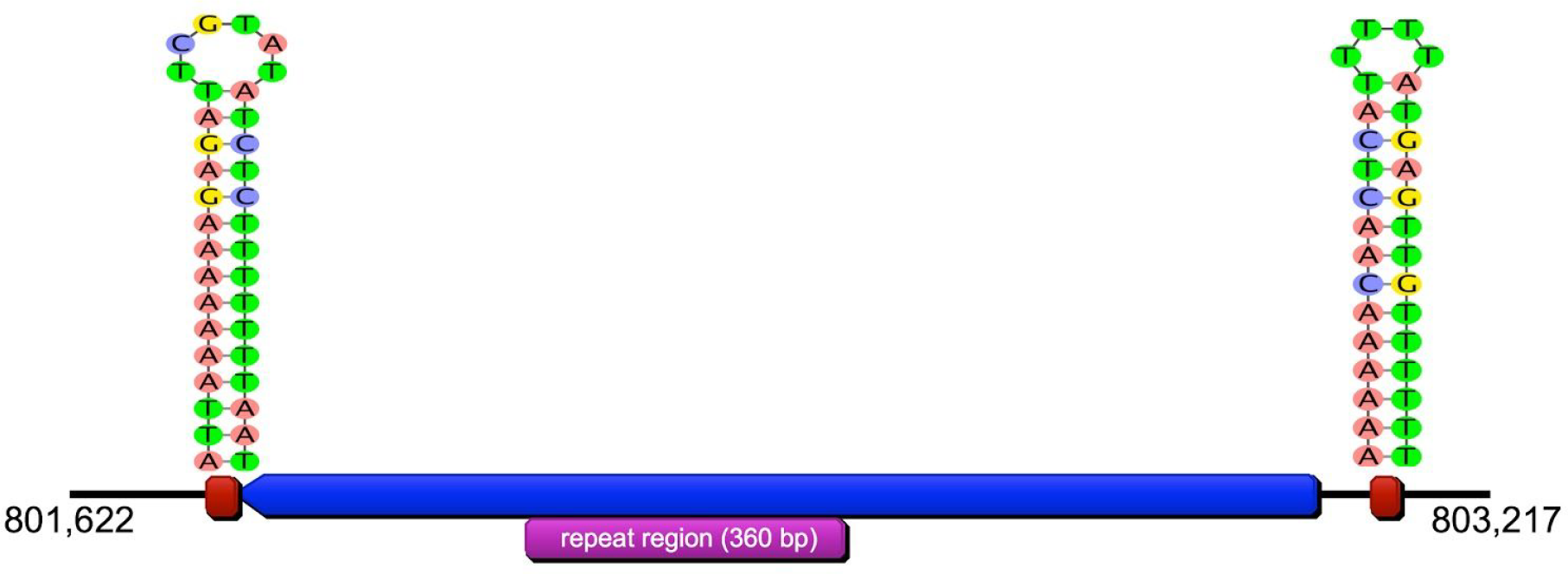
Position and structure of stem loops located in the UTR regions of the BD1-5 repeat protein. *Stemloop-3’* 5 bp from gene end to 31-bp downstream – ATTAAAAAAAGAGATTCGTATATCTCTTTTTTTAAT (ΔG = −10.75 kcal/mol); *stemloop-5’* 62-bp to 94-bp upstream – AAAAAACAACTCATTTTTATGAGTTGTTTTTT (ΔG =-12.49 kcal/mol). A highly similar (88% identical) stem-loop sequence was identified elsewhere in the genome within the 5’-end of a putative Type II/IV secretion system gene: AAAAAACACTGATAAAAATCAGTGTTTTTT (coordinates-167813-167842; ΔG = −11.49 kcal/mol).

**Figure S5:**
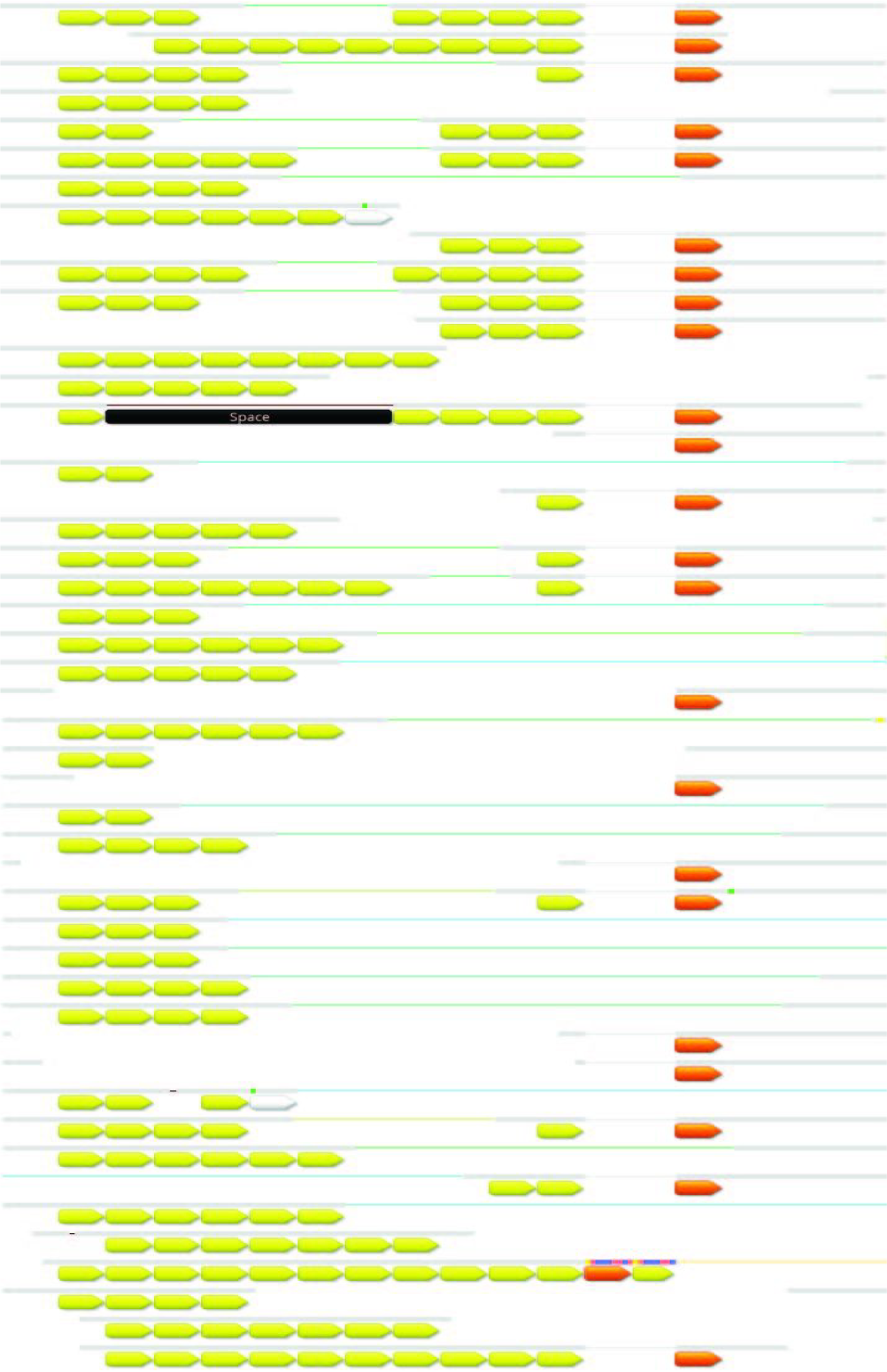
Small subset of the *Colwellia* gene predicted to include a PTD repeat. Repeats code for the tripeptide PDT. The black bar indicates a deletion in one variant. The sequence of yellow and orange 9-mers is provided in Figure 1.

**Additional files**: Newick version of the 16S rRNA tree with full sequence identifiers.

